## Supplemental Information for "In situ visualization of opioid and cannabinoid drug effects using phosphosite-specific GPCR antibodies"

**Supplementary information**

**This .pdf includes**

**Supplementary Figures: 2**

**Supplementary Tables: 1**

**Supplementary Figure 1**

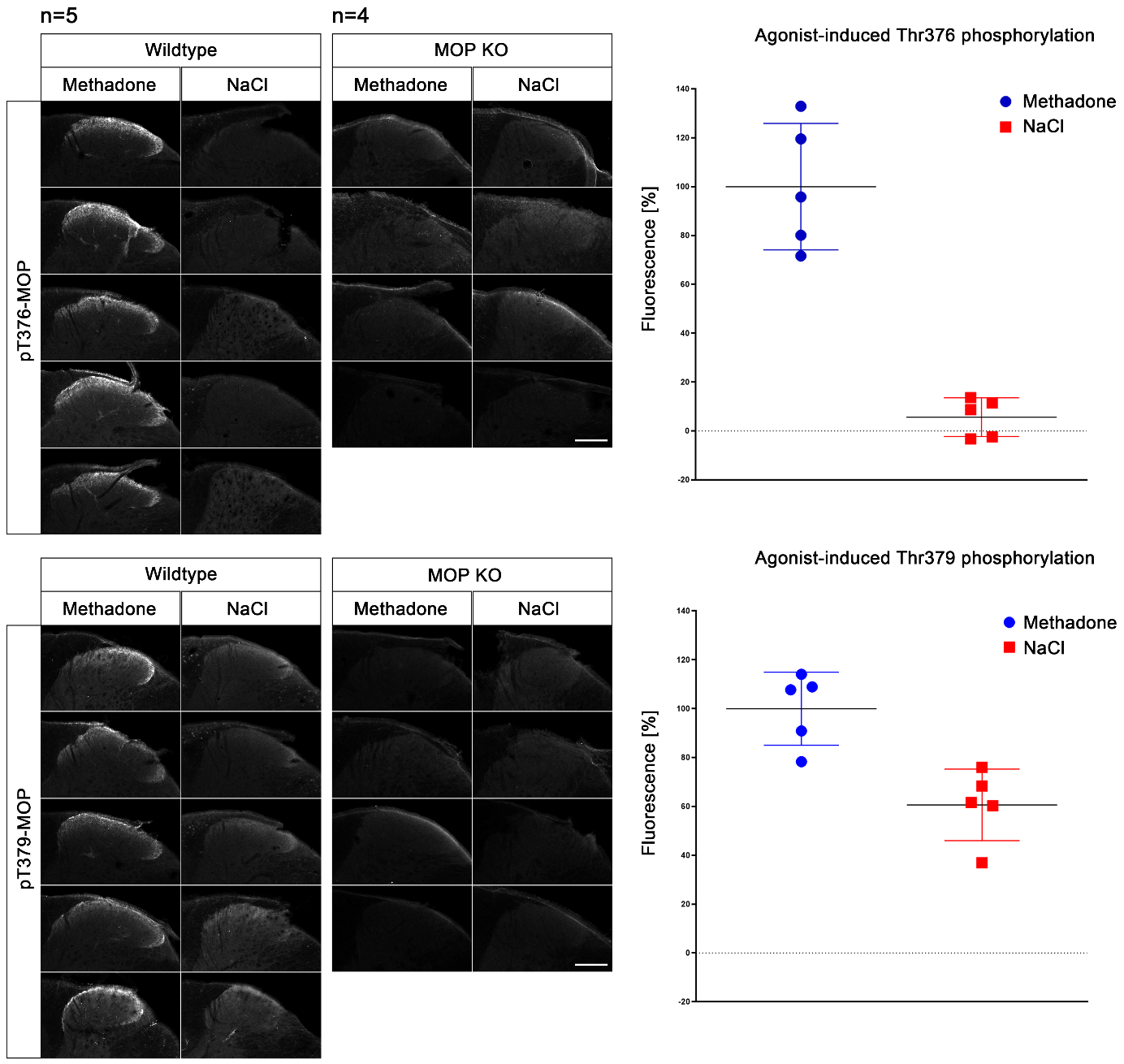
**Supplementary Figure 1. Comparison of phospho-MOP staining between different mice.** Animals were either treated with methadone or NaCl for 30 min, transcardially perfused, fixed and stained in the presence of PPIs. Shown are confocal images of coronal sections of the spinal cord of up to five different mice stained with either pT376-MOP or pT379-MOP antibody. MOP KO staining served as background control. Mean of five independent stained spinal cord slices is shown as 100%. Note that there is some range between the stained areas. pT379-MOP also shows a resilient phosphorylation in NaCl treated mice. ImageJ and GraphPad Prism were used for quantification. Scale bar: 250 µm.

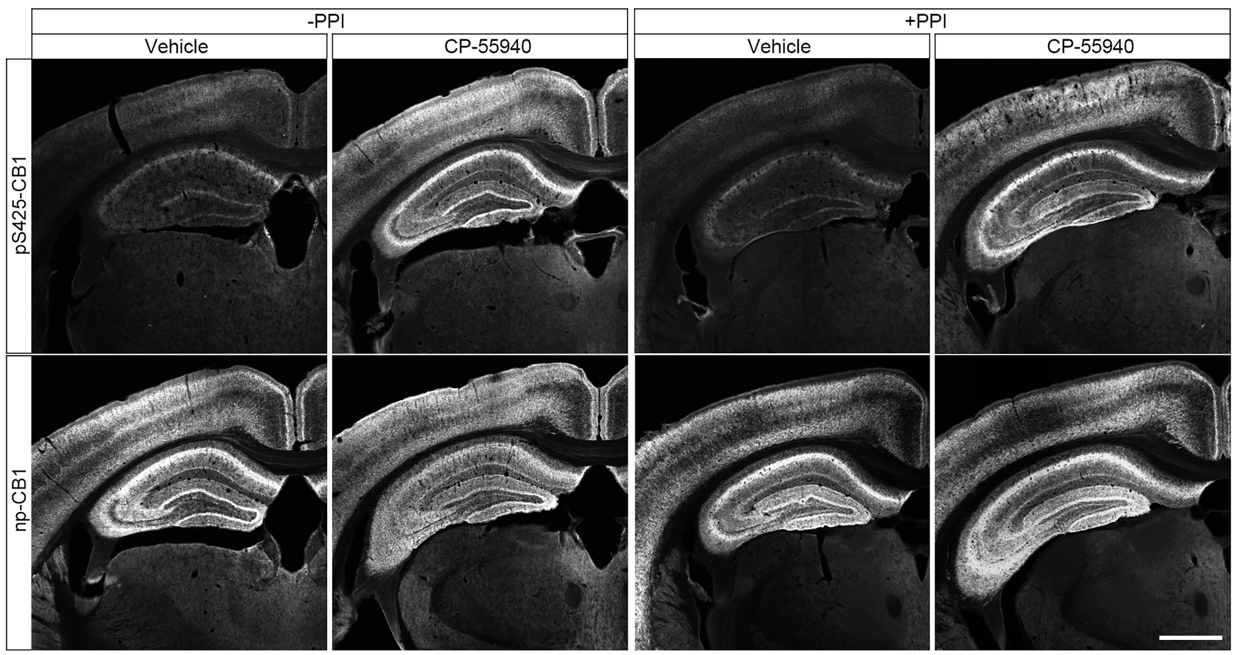

**Supplementary Figure 2. Comparison of phospho-CB1 immunohistochemistry in the presence or absence of protein phosphatase inhibitors.** Animals were either treated with CP-55940 or vehicle for 30 min, transcardially perfused, fixed and stained in the presence (+) or absence (-) of protein phosphatase inhibitors (PPI). Shown are confocal images of coronal sections of the brain stained with pS425-CB1 or np-CB1 antibody. Note that no PPIs are required to obtain agonist-induced phospho-CB1 immunostaining. Scale bar = 1000 µm.

**Supplementary Table 1:** Source of primary antibodies.

| **Antibody** | **Catalog#** | **Dilution** | **Vendor** |
| --- | --- | --- | --- |
| pT370-MOP | 7TM0319B | 1:500 | 7TM Antibodies |
| pS375-MOP | 7TM0319C | 1:100 | 7TM Antibodies |
| pT376-MOP | 7TM0319D | 1:400 | 7TM Antibodies |
| pT379-MOP | 7TM0319E | 1:400 | 7TM Antibodies |
| np-MOP | 7TM0319N | 1:4.000 | 7TM Antibodies |
| np-MOP | ab134054 | 1:500 | abcam |
| pS425-CB1 | 7TM0056A | 1:50 | 7TM Antibodies |
| np-CB1 | 1000659 | 1:3.500 | Cayman |
| np-CB1 | MFSR100610 | 1:4.000 | Frontier Institute |
